# Dysregulation of Sterol Regulatory Element-Binding Protein 2 Gene in HIV Treatment-Experienced Individuals

**DOI:** 10.1101/742486

**Authors:** Anuoluwapo Sopeyin, Lei Zhou, Min Li, Lydia Barakat, Elijah Paintsil

**Affiliations:** Department of Pediatrics, Yale University School of Medicine, New Haven, CT, USA; Department of Medicine, Yale University School of Medicine, New Haven, CT, USA; Department of Pharmacology, Yale University School of Medicine, New Haven, CT, USA; School of Public Health, Yale University, New Haven, CT, USA

**Keywords:** Antiretroviral therapy, toxicity, metabolic syndrome, cholesterol metabolism

## Abstract

We investigated the effects of antiretroviral therapy (ART) on cholesterol biosynthesis in a case-control study. mRNA and protein expressions of 3-hydroxy-3-methylglutaryl-coenzyme A reductase (HMGCR) and ATP binding cassette transporter A1 (ABCA1) were significantly upregulated in cases (HIV+) compared to controls (HIV-). We observed dysregulation between sterol regulatory element-binding protein 2 (SREBP-2, sensory control) and HMGCR and low-density lipoprotein receptor (LDLR) pathways. Dysregulation of cholesterol biosynthesis genes may predate clinical manifestation of ART-induced lipid abnormalities.

---

Antiretroviral therapy (ART)-associated metabolic derangement and metabolic syndrome (MetS) are more prevalent than ART-associated toxicities such as lactic acidosis, peripheral neuropathies, cardiomyopathies, and pancytopenia^1-5^. In adults, MetS is defined as having at least three out of five of the following components: impaired fasting glucose or diabetes, hypertension, central obesity (increased waist circumference), elevated triglycerides or reduced high-density lipoprotein (HDL) cholesterol^6^. The prevalence of MetS in people living with HIV (PLWH) is as high as 83%, particularly in PLWH on protease inhibitors (PI)-based regimens^7^, compared to 34% in the general population ^8^. MetS has been associated with an increased risk of cardiovascular diseases (CVDs) such as myocardial infarction (MI), atherosclerosis, and stroke ^9, 10^.

The high prevalence of MetS and CVDs in PLWH may be due to a complex interplay of HIV infection^11, 12^, ART exposure, other viral co-infections^13, 14^, and traditional risk factors such as genetic predisposition genetics^15^ and lifestyle habits. However, the underlying mechanisms are not well known. We recently observed that CEM cells exposed to 1x and 4x the C_max_ of various antiretroviral combinations resulted in differential expressions of 122 out of 48,226 genes using microarray analysis (published^16^ and unpublished data). Over a third of those genes belonged to the cholesterol biosynthesis pathway. Based on our findings, we hypothesized that ART could perturb cholesterol biosynthesis genes before manifestation of overt signs and symptoms of lipid abnormalities and MetS. We investigated the effect of ART on cholesterol biosynthesis in peripheral blood mononuclear cells (PBMCs) of HIV treatment-experienced individuals (cases) compared to HIV-negative healthy individuals (controls).

We interrogated four major pathways genes involved in cholesterol regulation using mRNA and protein expression studies: sensory control (sensor sterol regulatory element binding protein 2, SREBP-2), de novo synthesis (3-hydroxy-3-methylglutaryl-coenzyme A reductase, HMGCR), cholesterol uptake (low-density lipoprotein receptor, LDLR), and efflux (ATP binding cassette transporter A1, ABCA1). We also measured the expression of AMP-activated protein kinase A1 &B2 (AMPK A1 &AMPK B2, precursors of the cholesterol synthesis pathway.

Study participants were enrolled at the Yale-New Haven Hospital from April 2011 to March 2013. The details of the study design for the study cohort have been described previously^17^. In brief, cases comprised HIV-infected individuals on ART for at least 12 months without clinical and/or laboratory toxicities. Cases were matched HIV-negative controls. by age, sex, and race/ethnicity to HIV-negative controls. The study protocol was approved by the Institutional Review Board of the Yale School of Medicine. All participants gave their written informed consent before participation in the study.

The cholesterol sub-study, included only study participants with sufficient archived PBMCs for the analysis (cases, n=18, and controls, n=18). We extracted RNA and proteins from PBMCs for mRNA expression and protein expression, respectively, of cholesterol biosynthesis pathway genes. We report data as medians with interquartile ranges (unless otherwise stated) and as frequencies with percentages for continuous and categorical variables, respectively. We used Wilcoxon signed-rank test to compare continuous variables and Pearson correlation to examine bivariate associations. P-values were considered significant if <0.05. All statistical analyses were performed using GraphPad Prism software.

The demographic and clinical characteristics of study participants are illustrated in Table 1. The mean age was 52 years (range, 30 to 72 years), with 67% being males. The ethnicity comprised 28% Non-Hispanic whites, 6% Hispanic white and 66% African Americans. The ART regimen of the cohort was mostly tenofovir/emtricitabine (33%) plus PI or integrase inhibitor, tenofovir/emtricitabine/efavirenz (50%) and zidovudine/lamivudine (17%).

**Table 1.** Demographic and clinical characteristics of study participants

| Variable |  | HIV un-infected individuals<br>(Controls, n=18) | HIV infected individuals<br>on antiretroviral therapy<br>(Cases, n=18) |
| --- | --- | --- | --- |
| Mean Age (Range), years |  | 52 (32-72) | 52 (30-68) |
| Gender | Male | 12 | 12 |
|  | Female | 6 | 6 |
| Race | White non-Hispanic | 5 | 5 |
|  | White Hispanic | 1 | 1 |
|  | African American | 12 | 12 |
| Mean CD4 count (range)<br>(count/μL) | | N/A | 735 (264-1159) |
| Mean Viral load (range)<br>(copies/mL) |  | N/A | 23 (20-79) |
| Mean Duration of exposure to<br>treatment (range) (yrs) |  | N/A | 4.77 (1-7.5) |
| Treatment Regimen (%) | NRTI | N/A | 9 |
|  | NRTIs/NNRTI | N/A | 9 |
| Mean Cholesterol (range) |  | N/D | 175 (72-248) |
| Mean HDL (range) |  | N/D | 52 (25 -82) |
| Mean LDL (range) |  | N/D | 99 (9-158) |
| Mean Triglycerides (range) |  | N/D | 129 (56-258) |

N/A, not applicable
N/D, not determined for healthy volunteers

We quantified the mRNA expression of genes using qRT-PCR (*supplementary Table 1* for primer sets of the genes). Data was obtained from duplicate assays performed on at least two different occasions. The fold change in gene expression was calculated using the formula 2^Δ*CT*^ (Ref), where Δ*C*_*T*(case)_ = (*C*_*T*(gene of interest)_ − *C*_*T*(GAPDH)_), and Δ*C*_*T*(healthy control)_ = (*C*_*T*(gene of interest)_ − *C*_*T*(GAPDH)_). mRNA expressions of HMGCR and ABCA1 were upregulated in cases compared to controls (p=0.03 and p<0.01, respectively) (Figure 1). There was no significant difference in SREBP-2 expression among cases and controls, although cases tended to have a lower expression of SREBP-2 and AMPK B2 compared to controls (Figure 1).

**Figure 1.**
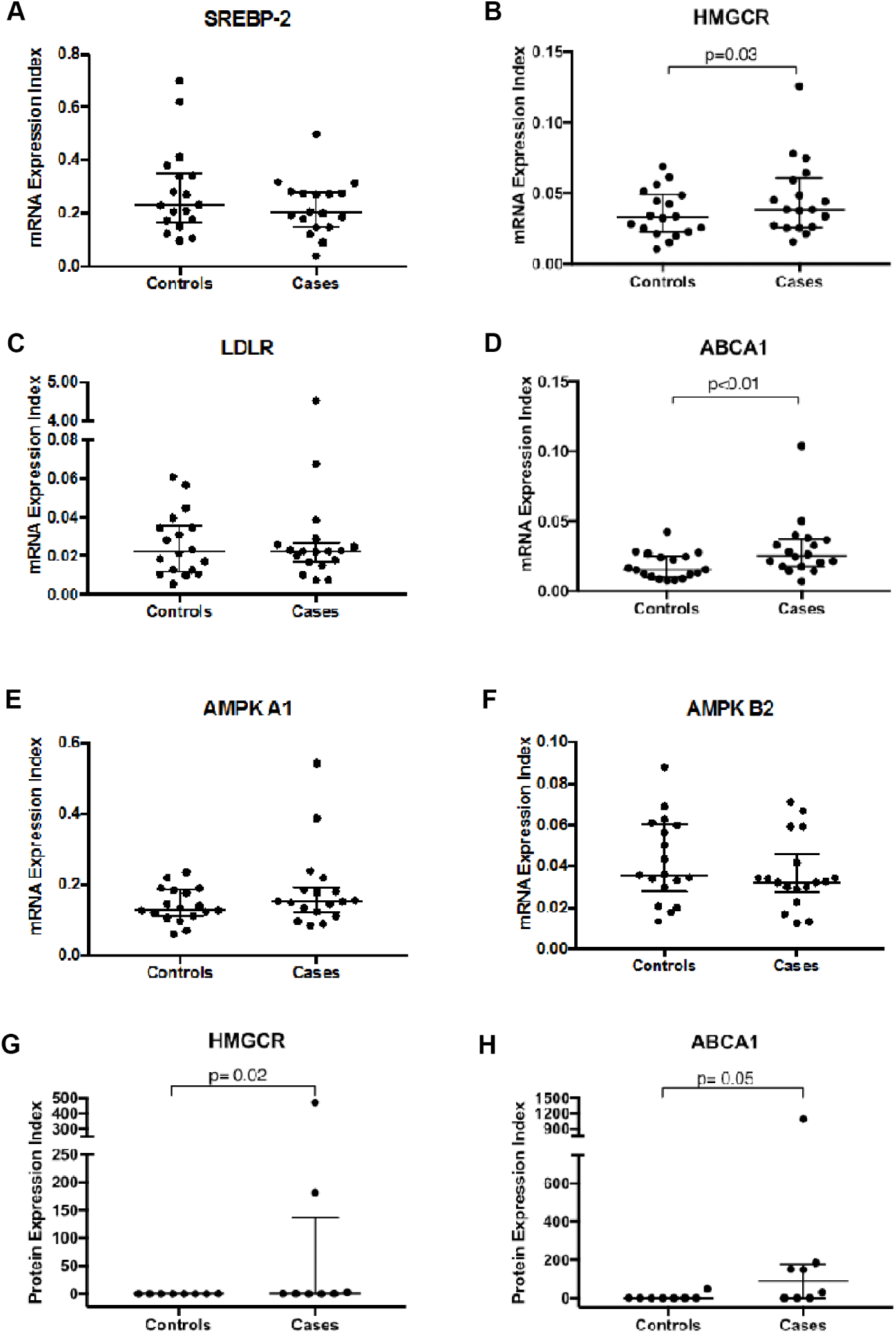
mRNA expression of cholesterol biosynthesis genes in peripheral blood mononuclear cells (PBMCs) of HIV positive individuals on ART (cases: N=18) and HIV negative individuals (controls: N=18). A. Sterol response element binding protein 2 (SREBP-2). B. HMG coenzyme reductase A (HMGCR). C. Low-density lipoprotein receptor (LDLR). D. Adenosine triphosphate–binding cassette transporter A1 (ABCA1). E. AMP Kinase A1 (AMPK A1). F. AMP Kinase A2 (AMPK A2). Protein expression of cholesterol biosynthesis genes in PBMCs of HIV positive individuals on ART (cases: N=8) and HIV negative individuals (controls: N=8). G. HMGCR. F. ABCA1.

With the significant increase in mRNA expressions of HMGCR and ABCA1 in cases, we investigated whether this translated to protein expressions. Western blot analysis was performed as described previously^18^ using tubulin as the housekeeping gene. The Western blot analysis was conducted on 16 participants with sufficient samples (cases n=8 and controls n=8). The density of the bands was quantified suing Quantity One Analysis Software. We observed a corresponding increase in protein expressions of HMGCR and ABCA1 (p=0.02, p=0.05, respectively) in cases (Figure 1G-H).

In health, there is a positive correlation between SREBP-2 and HMGCR as well as SREBP-2 and LDLR^19, 20^. Therefore, with differential upregulation of HMGCR and ABCA1 in cases and controls, we investigated the correlation between SREBP-2 and HMGCR, and SREBP-2 and LDLR. As expected in controls (N=18), there was a positive correlation between SREBP-2 and HMGCR (R^2^=0.24, p=0.04), and LDLR (R^2^=0.23, p=0.05) (Figure 3). To our surprise, the correlations among the cases were negative (Figure 3).

Under physiologic conditions, if intracellular levels of cholesterol become low, SREBP-2 is cleaved from the endoplasmic reticulum (ER) and migrates into the nucleus^21^ causing increased expression of HMGCR (the rate-limiting step in the synthesis of cholesterol) and LDLR (a membrane-associated cholesterol receptor)^22^ and decreased expression of ABCA1 (a cholesterol efflux protein). These regulatory mechanisms ensure cellular homeostasis. If there is an intracellular accumulation of cholesterol, the expression of ABCA1 is increased to facilitate reverse cholesterol transport out of the cell^23^. HIV infection disrupts cholesterol efflux by ABCA1, however, there is no consensus on the effect of ART on ABCA1 expression ^24, 25^.. ABCA1 is a member of a superfamily of ATP-binding cassette (ABC) transporters that are involved in transporting molecules across cellular membranes. ABCA1 is involved with exporting phospholipids and cholesterol to apolipoproteins to form HDL cholesterol. Mutations in ABCA1 have been associated with low levels of HDL as observed in Tangier Disease^26, 27^. There are reports of HIV treatment-naïve individuals with upregulation of ABCA1, which normalizes upon initiation of ART ^24^. Other studies have found downregulation of ABCA1 in HIV treatment naïve individuals^28^. The downregulation of ABCA1 in HIV treatment-naïve individuals has been implicated on HIV Nef protein; Nef protein downregulates ABCA1 leading to impaired efflux of cholesterol resulting in intracellular accumulation of cholesterol ^29-31^. This effect is reversed with initiation of ART^32^. The mean viral load of our cases was 23 (range, 20-79) copies/mL (Table 1). Therefore, the effect of HIV Nef protein may not play a significant role in our cohort. We expected that ART will normalize expression of ABCA1, however, there was upregulation of ABCA1 in cases. The mean duration of therapy among cases was 4.77 (range, 1-7.5) years. Is it plausible that the continued exposure to ART in our cohort led to upregulation of ABCA1?

It is also interesting that we found an upregulation in the expressions of HMGCR mRNA and protein; this is counterintuitive with upregulation of ABCA1. A plausible explanation is that the upregulation of HMGCR was the inciting event that resulted in intracellular cholesterol accumulation. Thus, the cell, in an attempt to restore homeostasis, increases the expression of ABCA1. Another possible explanation could be that there is dysregulation of cholesterol biosynthesis. We cannot tease out this conundrum without intracellular cholesterol levels. However, the serum levels of cholesterol in cases was within normal range implying the increase in ABCA1 is not translating to increased efflux from the cell (Table 1).

In the absence of intracellular cholesterol data, did a correlation analysis of mRNA expression of cholesterol biosynthesis genes. We observed a normal association of genes involved with cholesterol sensing—SREBP-2, HMGCR and LDLR in controls (Figure 2). In cases, there was negative correlation between SREBP-2 and HMGCR or LDLR (Figure 2). In health, SREBP-2 senses intracellular cholesterol levels and upregulates cholesterol synthesis via HMGCR and uptake from the extracellular environment via LDL receptors (LDLR)^33, 34^. Our finding suggests a dysregulation of cholesterol biosynthesis, particularly sensing by SREBP-2 in cases.

**Figure 2.**
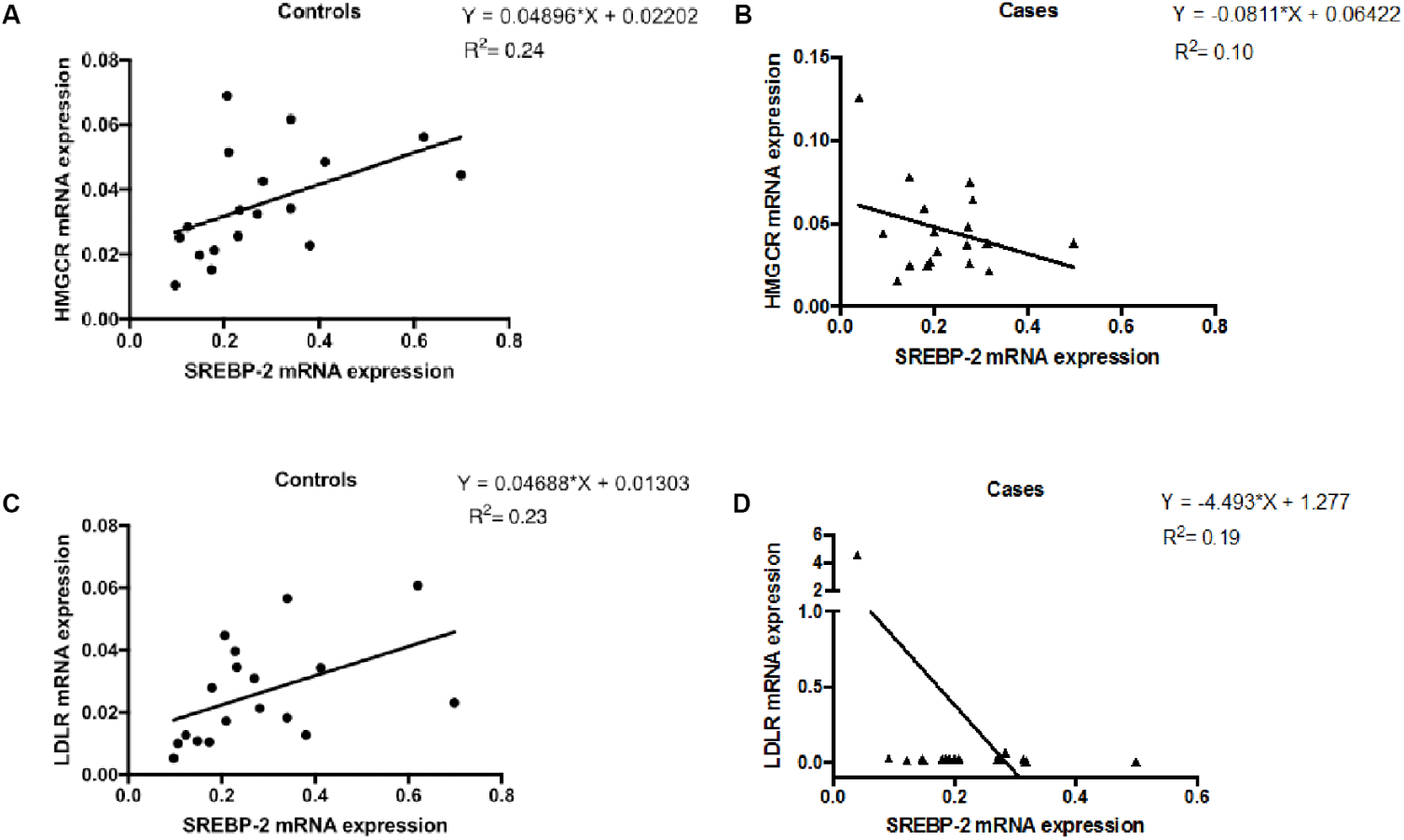
Correlation analysis of mRNA expression of cholesterol biosynthesis genes in peripheral blood mononuclear cells (PBMCs) of HIV positive individuals on ART (cases: N=18) and HIV negative individuals (controls: N=18). A. The mRNA expression of SREBP-2 vs. HMGCR in controls. B. The mRNA expression of SREBP-2 vs. HMGCR in cases. C. The mRNA expression of SREBP-2 vs. LDLR in controls. D. The mRNA expression of SREBP-2 vs. LDLR in cases.

Although, our study is one of the first studies to report potential dysregulation of cholesterol biosynthesis in HIV treatment-experienced individuals, it has several limitations. First, it is a cross-sectional study and not designed to assess causality. Second, it was an exploratory pilot sub-study with small sample size to test and generate hypotheses. Third, we did not quantify intracellular cholesterol levels to assess the effect of intracellular cholesterol on the genes studied. Fourth, the effect of HIV infection itself, say through HIV Nef protein, was not assessed since we did not have access to HIV treatment-naïve individuals with higher viral loads. Further studies are need with larger sample size and design to validate our findings. If our findings are validated, cholesterol biosynthesis genes could serve as biomarkers for predicting PLWH who will develop MetS and/or other lipid abnormalities and also for monitoring of treatment response of MetS and/or other lipid abnormalities in PLWH.

## Acknowledgements

This study was supported by grants from the National Institutes of Health (K08AI074404 to E.P.). A.S. was supported by an IDSA Medical Scholar Award.

